# MHC-matched allogeneic bone marrow transplants fail to eliminate SHIV-infected cells from ART-suppressed Mauritian cynomolgus macaques

**DOI:** 10.1101/2021.04.16.440168

**Authors:** Jason T. Weinfurter, Saritha S. D’souza, Lea M. Matschke, Sarah Bennett, Laurel E. Kelnhofer-Millevolte, Kran Suknuntha, Akhilesh Kumar, Jennifer Coonen, Christian M. Capitini, Peiman Hematti, Thaddeus G. Golos, Igor I. Slukvin, Matthew R. Reynolds

## Abstract

**Background:** Allogeneic hematopoietic stem cell transplants (allo-HSCTs) dramatically reduce HIV reservoirs in antiretroviral therapy (ART) suppressed individuals. However, the mechanism(s) responsible for these post-transplant viral reservoir declines are not fully understood but may include pre-transplant conditioning regimens, ART-mediated protection of donor cells, and graft-versus-host (GvH) responses. Therefore, we modeled allo-HSCT in ART-suppressed simian-human immunodeficiency virus (SHIV)-infected Mauritian cynomolgus macaques (MCMs) to illuminate factors contributing to transplant-induced viral reservoir decay.

**Results:** We infected four MCMs with CCR5-tropic SHIV162P3 and started ART 6-16 weeks post-infection (p.i.) to establish robust viral reservoirs. We maintained the MCMs on continuous ART during myeloablative conditioning with total body irradiation (TBI) and while transplanting allogeneic MHC-matched α/β T cell-depleted bone marrow cells. Post-transplant, we prophylactically treated the MCMs with cyclophosphamide and tacrolimus to prevent GvH disease (GvHD). The transplants produced ~85% whole blood donor chimerism without causing high-grade GvHD. Consequently, three MCMs had undetectable SHIV DNA in their peripheral blood mononuclear cells post-transplant. However, SHIV-harboring cells persisted in various tissues. We detected viral DNA in lymph node biopsies and terminal analyses of tissues between 38 and 62 days post-transplant. Further, we removed ART from one MCM at 63 days post-transplant, resulting in viral rebound within seven days and viral loads nearing 1×10^8^ SHIV RNA copies/ml of plasma after treatment interruption.

**Conclusions:** Our results indicate that myeloablative conditioning and maintaining ART through the peri-transplant period alone are insufficient for eradicating latent viral reservoirs early after allo-HSCTs. Furthermore, our findings suggest that extended ART and GvH responses may be necessary to substantially deplete viral reservoirs after allo-HSCTs.

## Background

Allogeneic hematopoietic stem cell transplants (allo-HSCTs) are essential curative therapies for high-risk hematologic malignancies [1–3], replacing recipient cells of hematopoietic origin with donor cells. In persons with HIV, allo-HSCTs also dramatically reduce viral reservoirs, enabling several transplant recipients to prevent or delay viral rebound after stopping antiretroviral therapy (ART) [4–10]. While allo-HSCTs are potent cellular immunotherapies, they are not scalable nor relevant to most people living with HIV without underlying hematologic cancers. Therefore, understanding the mechanism(s) of allo-HSCT-mediated reservoir decay can aid in designing safer and more efficacious treatment regimens. To this end, allo-HSCT components postulated to reduce viral reservoirs include pre-transplant conditioning regimens, protecting donor CD4^+^ T cells from infection with ART, graft-versus-host (GvH) responses, and donor HSCs lacking functional C-C chemokine receptor type 5 (CCR5) [11, 12].

To enter cells, HIV must sequentially bind to CD4 and a cellular coreceptor, primarily chemokine receptors CCR5 and CXCR4. These molecular interactions are disrupted in individuals homozygous for the *CCR5*Δ*32* gene variant (Δ*32*/Δ*32*), which prevents functional CCR5 from being expressed on cell surfaces, making Δ*32*/Δ*32* cells resistant to infection with CCR5-tropic HIV strains. The protective properties of Δ*32*/Δ*32* cells were employed in the hematologic cancer treatments of two men with HIV. In these men, transplanting Δ*32*/Δ*32* HSCs induced cancer remission and the rapid depletion of HIV reservoirs, allowing the “Berlin” and “London” patients to stop ART for at least four years without viral rebound [5, 6, 8, 13]. Undoubtedly, the absence of CCR5 on donor cells prevented virus replication and the reestablishment of viral reservoirs post-transplant, supporting long-term HIV remission. However, additional transplant-related factors must have cleared pre-existing viral reservoirs.

Transplants with Δ*32* heterozygous or CCR5 wild-type donors in ART-suppressed recipients provide further insights into eliminating latent HIV [4, 7, 9, 14, 15]. Notably, three CCR5 wild-type HSC recipients substantially reduced their viral reservoirs post-transplant and delayed HIV rebound up to 9 months after analytical treatment interruption (ATI) [4, 15]. Mathematical modeling suggests that only hundreds to thousands of latently infected cells remained in these individuals at ART withdrawal [16]. Assuredly, maintaining ART during the peri-transplant period was an essential component of the allo-HSCT regimens, limiting HIV’s spread from recipient cells to vulnerable donor cells [17]. Importantly, these allo-HSCTs also indicate that factors outside of donor genetics contribute to depleting viral reservoirs.

An attractive explanation for allo-HSCT-induced reservoir decay is the pre-transplant conditioning regimens used to reduce malignant cell burdens, create space for HSC engraftment, and ablate immune cells mediating graft rejections [11, 18]. However, studies in humans and nonhuman primates (NHPs) indicate that pre-transplant conditioning regimens only transiently reduce viral reservoirs [19–24], which are refilled by the homeostatic or antigen-driven proliferation of latently infected cells [17, 22]. Thus, cytoreductive treatments alone are insufficient for eradicating HIV reservoirs, even when ART is maintained.

A more likely mechanism for allo-HSCT-mediated viral reservoir decay is the beneficial effects of GvH responses [7, 15, 25]. After allo-HSCTs, major histocompatibility complex (MHC)- matched donor T cells recognize minor alloantigens expressed by host leukocytes, eliminating malignant and HIV-infected cells simultaneously, thereby mediating graft-versus-tumor and graft-versus-viral reservoir (GvVR) effects, respectively [26, 27]. But these GvH interactions can also result in life-threatening graft-versus-host disease (GvHD), a condition where donor cells attack non-hematopoietic origin cells or tissues. Disentangling allo-HSCT’s GvVR effects from GvHD may improve the safety and efficacy of cellular therapies targeting viral reservoirs.

In humans, variability in treatment regimens makes it challenging to investigate allo-HSCTs effects on HIV reservoirs. However, allo-HSCT treatment conditions can be standardized in NHP models of HIV infection. To this end, cynomolgus macaques (*Macaca fascicularis*) from the island of Mauritius are well-suited to model allo-HSCTs in people with HIV. Mauritian cynomolgus macaques (MCM) are descended from a small founder population [28], resulting in limited genetic diversity, even at polymorphic loci. As a result, MCMs have only seven MHC haplotypes (termed M1-M7), facilitating the assembly of MHC-matched transplant pairs [29–31]. Moreover, SIV and simian-human immunodeficiency viruses (SHIVs) infections in MCMs recapitulate HIV replication and viral reservoir dynamics [30–33], providing a platform for studying allo-HSCTs impact on viral reservoirs.

In this study, we modeled CCR5 wild-type allo-HSCTs in four SHIV-infected and ART-suppressed MCMs using our recently established TCRα/β-depleted MHC-matched bone marrow (BM) transplant model, which supports donor chimerism without GvHD [34]. As anticipated, the myeloablative conditioning regimen substantially depleted circulating leukocytes. As a result, we detected approximately 85% whole blood donor chimerism, with three MCMs having undetectable SHIV DNA in their peripheral blood mononuclear cells (PBMCs) post-transplant. However, SHIV-harboring cells persisted in various tissues, indicating that ART-suppression, allo-HSCs, and cytoreductive conditioning alone do not eliminate virally infected cells early after allo-HSCTs. Our findings suggest that extended ART treatments and GvH responses may be needed to eradicate viral reservoirs after allo-HSCTs.

## Results

### SHIV infection and ART suppression

We inoculated four MCMs (MCMs A-D) intravenously with SHIV162P3, a chimeric virus with a CCR5-tropic HIV envelope [35]. As anticipated, virus replication peaked by three weeks post-infection (p.i.) and declined thereafter (Fig. 1). To establish robust viral reservoirs, we allowed the SHIV infections to progress untreated for between 6-16 weeks p.i. before starting the MCMs on an ART regimen consisting of tenofovir, emtricitabine, and raltegravir. The ART regimen suppressed plasma viremia to undetectable levels within three weeks of initiation (<100 viral RNA (vRNA) copy/eq per ml; Fig. 1). We maintained ART for the duration of the study, except where noted below, keeping plasma viremia undetectable, save for single viral blips in MCMs A and D.

**Figure 1.**
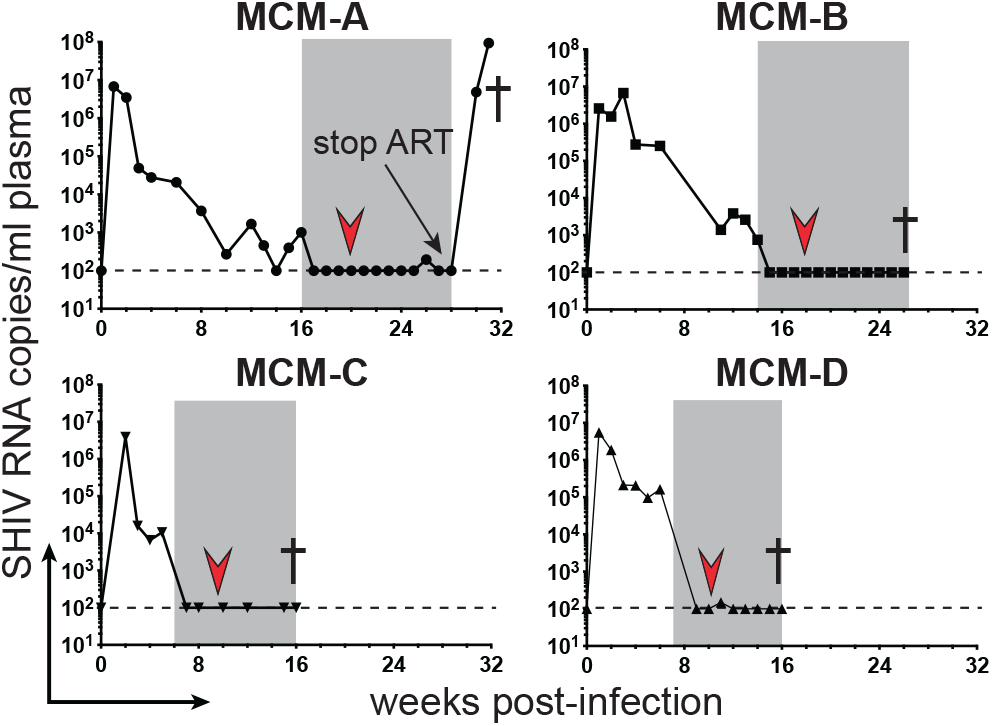
Plasma viral loads after SHIV162P3 infection. Longitudinal plasma viral loads for the SHIVI62P3-infected MCMs, with dashed lines indicating the qRT-PCR limit of detection (100 vRNA copy Eq/ml plasma). The gray boxes signify the timing of ART, the red arrows designate allo-HSCs infusions, and † indicate necropsies.

### Allo-HSCT in SHIV-infected, ART-suppressed MCM

Our goal was to model allo-HSCTs with CCR5 wild-type cells in ART-suppressed people with HIV while minimizing the likelihood of severe GvHD. We collected BM aspirates from sex and MHC-matched MCM (Table 1) and depleted cells expressing α/β T-cell receptors (TCRs), removing potentially alloreactive T cells [34]. Approximately one month after starting ART, we delivered myeloablative doses of total body irradiation (TBI; two consecutive days of 5 Gray (Gy), 10 Gy total) and infused the MCMs with α/β T cell-depleted HSCs (0.6-1.0×10^7^/Kg; Table 1). Post-transplant, we prophylactically treated the MCMs with cyclophosphamide and tacrolimus to prevent GvHD (Supplementary Table S1).

**Table 1.** MCM characteristics and doses of transplanted cells

| Table 1. MCM characteristics and doses of transplanted cells |  |  |  |  |  |  |  |  |  |  |  |  |  |
| --- | --- | --- | --- | --- | --- | --- | --- | --- | --- | --- | --- | --- | --- |
| RECIPIENT |  |  |  | DONOR |  |  |  | TOTAL CELLS TRANSPLANTED |  | CD34<sup>+</sup> CELLS |  | TCRα/β<sup>+</sup> CELLS |  |
| ID | MHC type | Sex | Age (yrs) | ID | MHC type | Sex | Age (yrs) | Total | Dose/Kg | Total | Dose/Kg | Total | Dose/Kg |
| MCM-A | M1/M2 | F | 4 | MCM-E | M2/M2 | F | 9 | 4.4 X10<sup>8</sup> | 1.5 X10<sup>8</sup> | 2.5 X10<sup>7</sup> | 0.8X10<sup>7</sup> | 3 X10<sup>5</sup> | 1 X10<sup>5</sup> |
| MCM-B | M1/M2 | F | 4 | MCM-F | M1/M2 | F | 3 | 8.5 X10<sup>8</sup> | 1.8 X10<sup>8</sup> | 5 X10<sup>7</sup> | 1 X10<sup>7</sup> | 17 X10<sup>5</sup> | 3.6X10<sup>5</sup> |
| MCM-C | M1/M2 | F | 13 | MCM-G | M1/M2 | F | 11 | 7.1 X10<sup>8</sup> | 1.2X10<sup>8</sup> | 4.8 X10<sup>7</sup> | 0.8 X10<sup>7</sup> | 7X10<sup>5</sup> | 1.1X10<sup>5</sup> |
| MCM-D | M1/M2 | M | 7 | MCM-H | M1/M2 | M | 6 | 8.6 x10<sup>8</sup> | 1.6 x10<sup>8</sup> | 3.4 x10<sup>7</sup> | 0.6 x10<sup>7</sup> | 8 x10<sup>5</sup> | 1.4 x10<sup>5</sup> |

As expected, the myeloablative conditioning regimen rapidly depleted peripheral white blood cells (WBCs; Fig. 2a). However, the MCMs also developed chronic thrombocytopenia post-transplant (Fig. 2b), which we treated with irradiated whole blood transfusions from SHIV-negative MCMs. Nevertheless, neutrophils reappeared in the peripheral blood of the MCMs by 21 days post-transplant, with absolute counts in MCM-D spiking above pre-transplant levels before rapidly declining (Fig. 2c). The remaining three MCMs had persistent neutropenia after their initial bursts of circulating neutrophils. Likewise, monocytes reemerged in the peripheral blood by 21 days post-transplant, with MCM-D again having the most robust recovery (Fig. 2d). The absolute monocyte counts were more variable than the neutrophil counts post-transplant, gradually declining in MCMs A and C, while rapidly increasing in MCMs B and D before their deaths.

**Figure 2.**
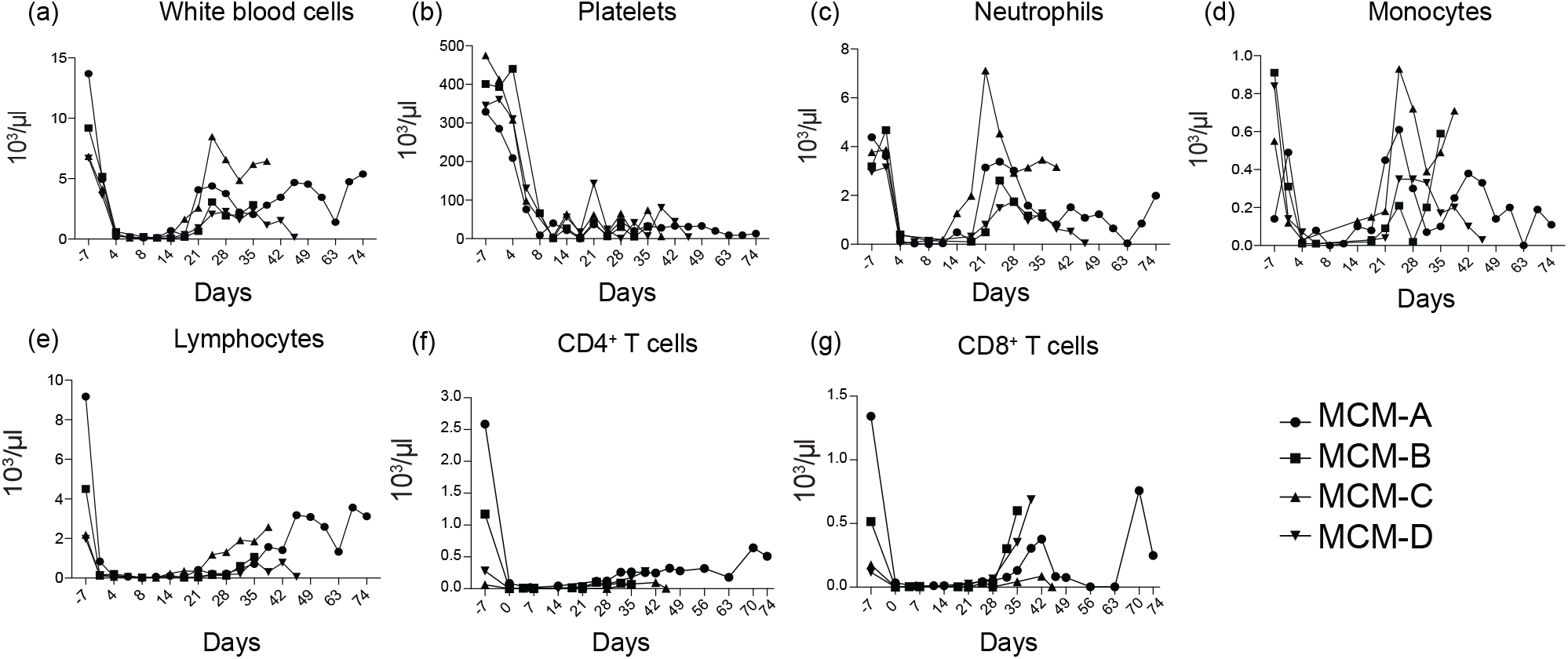
Effect of allo-HSCTs on cell populations and platelets. Longitudinal absolute counts of (a) white blood cells, (b) platelets, (c) neutrophils, (d) monocytes, (e) lymphocytes, (f) CD4^+^ T cells, and (e) CD8^+^ T cells after allo-HSCTs. Counts are displayed as x103/μl.

Lymphocytes recovered slowly after the allo-HSCTs (Fig. 2e). In MCM-D, lymphocytes reappeared in the peripheral blood approximately three weeks post-transplant and returned to near pre-transplant levels by day 38 post-transplant. Likewise, MCM-A exhibited a slow but steady increase in peripheral lymphocytes, reaching ~3,500 cells/μl by day 71 post-transplant. In contrast, MCMs B and C had stunted lymphocyte recoveries, with absolute lymphocyte counts failing to reach 1,000 cells/μl of blood by 5- and 7-weeks post-transplant, respectively. Similarly, CD4^+^ T cells failed to fully recover in the MCMs blood post-transplant (Fig. 2f). Finally, circulating CD8^+^ T cells increased in MCMs A, B, and D approximately one-month posttransplant (Fig. 2g), fluctuating in MCM-A until its death. Meanwhile, CD8+ T cells poorly recovered in MCM-C, only reaching 86 CD8+ T cells/μl blood at 42 days post-transplant.

We measured post-transplant donor engraftment by deep sequencing single-nucleotide polymorphisms (SNPs) unique to the donor and recipient MCMs. Despite low absolute leukocyte counts, the allo-HSCT recipients exhibited greater than 85% whole blood donor chimerism by three weeks post-transplant, followed by modest declines to approximately 80% donor chimerism (Fig. 3a). Notably, MCM-A, the longest surviving transplant recipient, maintained 70-90% donor chimerism for two months until euthanasia. Additionally, we measured donor chimerism of circulating myeloid (monocytes) and lymphoid (B and T cells) lineages in MCMs A, C, and D at necropsy (cells from MCM-B were unavailable for analysis). The frequencies of donor-derived myeloid and lymphoid cells in MCMs C and D ranged between 65-93% of total cells (Fig. 3b), consistent with the whole blood chimerism results (Fig. 3a). In MCM-A, 78% of the B and T cells were of donor origin (Fig. 3b), similar to the 83% of donor cells detected in the whole blood. However, only 45% of the monocytes were of donor origin. Collectively, these results indicate that donor leukocytes replaced recipient cells early after transplant, with notable donor lymphocyte engraftment despite using α/β TCR-depleted grafts.

**Figure 3.**
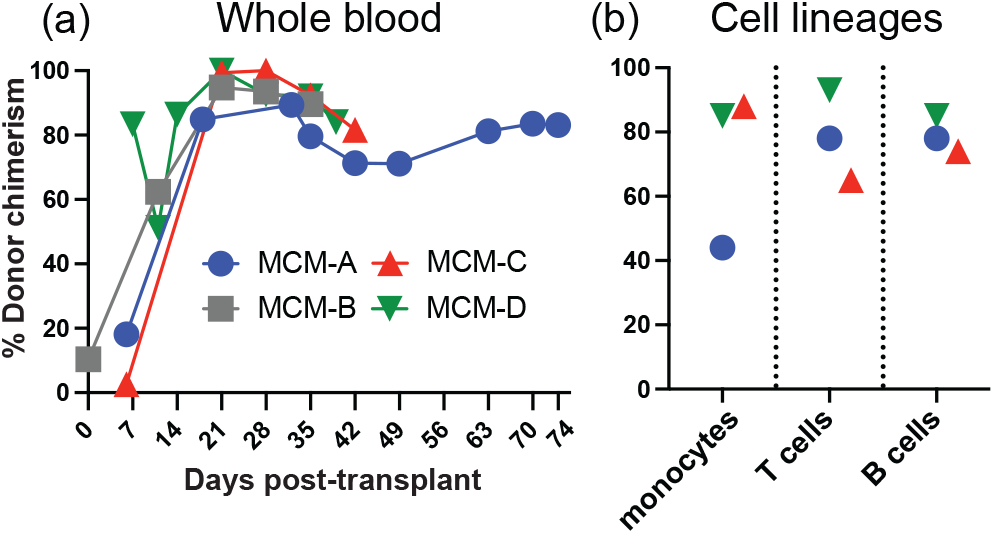
Donor chimerism after allo-HSCT. We determined donor chimerism levels post-transplant by sequencing SNPs differing between the donor and recipient. (a) Longitudinal percent donor chimerism in whole blood after allo-HSCT. (b) At necropsy, the percent donor chimerism among flow cytometry sorted PBMC monocytes (CD45^mid^CD14^+^ lymphocytes), T cells (CD14^-^CD45^hi^CD3^+^ lymphocytes), and B cells (CD14^-^CD45^hi^CD^20+^).

### Pre- and post-ART viral reservoirs

We designed our GvHD prophylaxis regimen of cyclophosphamide and tacrolimus to minimize high-grade GvHD. As anticipated, GvHD was limited post-transplant with only MCM-A exhibiting transient skin lesions (white flaking/dry skin) that resolved with minimal intervention. Additionally, MCMs C and D had diarrhea, which may have been a TBI side effect. Furthermore, the MCMs had no significant abnormalities in their liver functions or elevated total bilirubin posttransplant, indicating an absence of hepatic GvHD.

We measured cell-associated vDNA in PBMCs and inguinal lymph nodes to determine the effect of ART, myeloablative conditioning, and allo-HSCTs on viral reservoirs in the absence of severe GvHD (Fig. 4). Before starting ART, we detected more SHIV-harboring cells in the inguinal lymph nodes (range 743-2,680 copy Eq/1×10^6 cells) than the PBMC (range 255-512 copy Eq/1×10^6 cells), consistent with lymph nodes being a primary viral reservoir [36, 37]. Additionally, the lymph nodes of MCMs A and B contained more SHIV-infected cells than MCMD, potentially due to these MCMs starting ART later in infection (14-16 weeks versus 6 weeks). Lastly, the number of SHIV-infected cells declined in the blood of all the MCMs between 36-64 days post-transplant, with MCMs A, B, and D having undetectable vDNA in their PBMCs (Fig. 4a).

**Figure 4.**
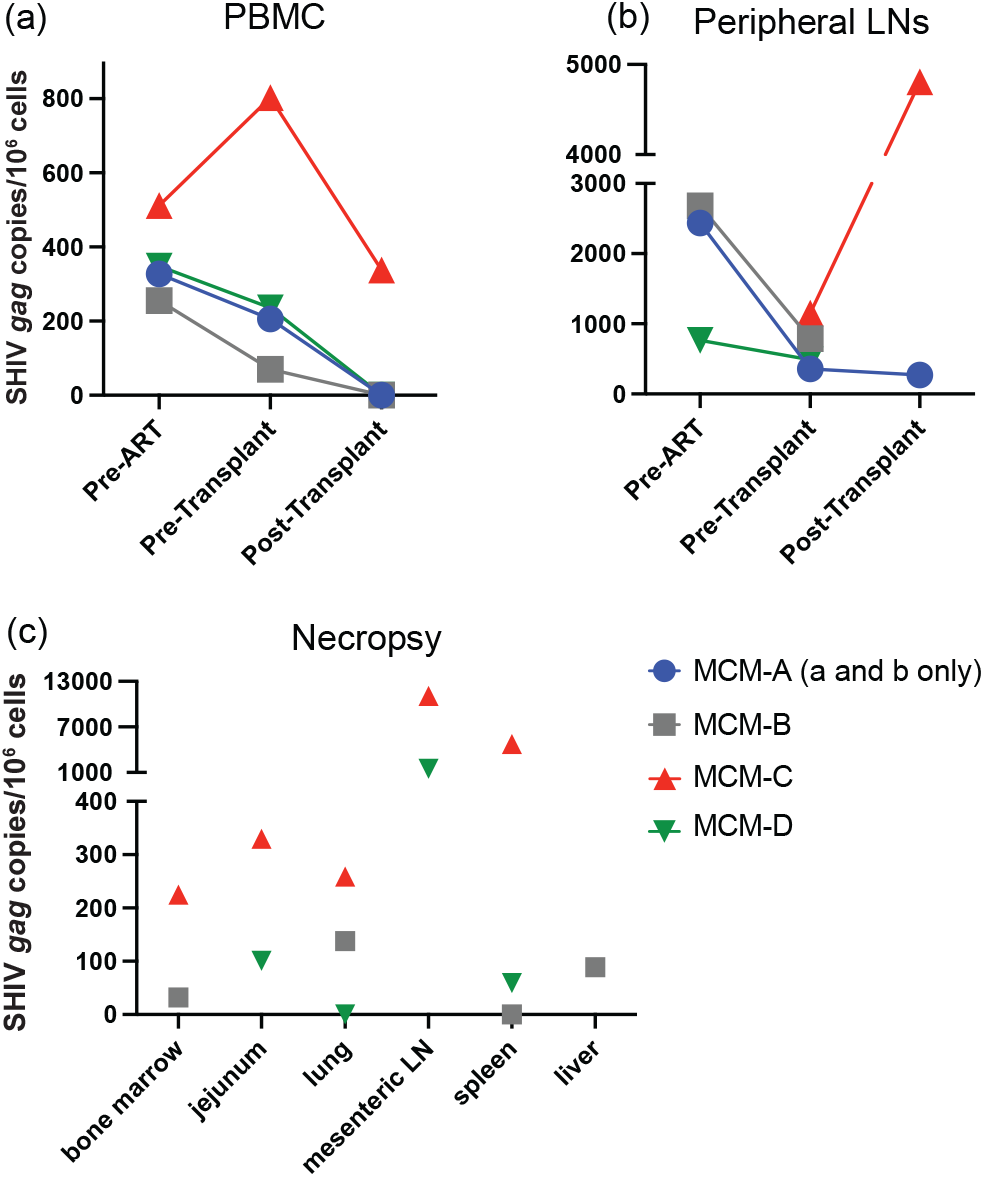
SHIV DNA in allo-HSC transplanted MCMs. We measured copies of vDNA in the (a) PBMC and (b) peripheral lymph nodes before ART initiation, after ART but before allo-HSCTs, and post-transplant. (c) SHIV DNA levels in the tissues of MCMs B, C, and D at necropsy. Total SHIV DNA was measured by digital droplet PCR using SIV gag-specific primers and normalized to 10^7^ cells by measuring macaque RNaseP p30, RPP30, in companion ddPCR assays.

Nevertheless, virally infected cells persisted in other tissues. MCMs-B, -C, and -D remained on ART until being necropsied, enabling a more thorough examination of posttransplant viral reservoirs. In MCM-C, we detected more vDNA in the inguinal lymph nodes at necropsy than in pre-transplant samples (1,151 vs. 4,807 vDNA copy Eq/million cells; Fig. 4b). This animal developed viral-induced colitis with ulcers post-transplant, potentially expanding latently infected cells or promoting localized virus replication, spreading SHIV-harboring cells into secondary lymphoid tissues. Nevertheless, we found vDNA outside of the secondary lymphoid tissues in the three MCMs, including the bone marrow, lung, and jejunum (Fig. 4c). Together these results demonstrate that viral reservoirs persist in various tissues early after allo-HSCTs without GvHD.

### SHIV rapidly rebounds after removing ART

ATIs are used in HIV cure studies to evaluate treatment efficacy [38, 39]. Therefore, we modeled ATI in MCM-A, an animal with a single viral blip but otherwise suppressed plasma viremia while on ART. One day before stopping ART, day 62 post-transplant, we collected blood and axillary lymph nodes to measure cell-associated SHIV. In these samples, we found no vDNA in the PBMCs but detected 271 vDNA copies/million cells in the lymph nodes, a slight decrease from pre-transplant inguinal lymph nodes (359 vDNA copies/million cells; Fig. 4b). Nevertheless, SHIV rebounded rapidly after removing ART (Fig. 1). Plasma viremia was 1×10^7^ vRNA copy eq/ml seven days after stopping treatment, roughly equivalent to peak acute viremia, and spiked to nearly 1×10^8^ vRNA copy eq/ml plasma at 10 days post-ART. These results show that latent reservoirs can reactivate shortly after treatment interruption and initiate viral rebound early after allo-HSCTs.

## Discussion

Allo-HSCT-associated HIV reservoir decay likely results from parallel processes protecting donor cells from infection and eliminating pre-existing latently infected cells [10, 11]. Here, we employed the clinically relevant SHIV/MCM model to determine the effect of continuous ART and allo-HSCTs on viral reservoirs in the absence of high-grade GvHD. We recently reported that transplanting MCMs with MHC-matched TCRα/β^+^-depleted HSCs and prophylactically treating them with cyclophosphamide and tacrolimus prevents GvHD posttransplant [34]. Employing this regimen, only one SHIV-infected MCM had transient skin lesions post-transplant. These results contrast with previous NHP allo-HSCT studies where T-cell replete HSCs quickly induced GvHD in ART-suppressed SHIV-infected rhesus macaques [40] and uninfected MCMs [41].

Although we prevented GvHD, removing alloreactive TCRα/β+ T cells from the bone marrow grafts may have inhibited viral reservoir clearance. Still, our results agree with a previous study transplanting MHC-haploidentical HSCs into ART-suppressed SHIV-infected rhesus macaques that developed GvHD post-transplant [49]. Both studies observed reduced SHIV DNA levels in the blood and lymph nodes post-transplant, with SHIV reservoirs persisting in other tissues. Further, post-transplant follow-up was short in both studies, supporting the need for extended times on ART post-transplant to clear pre-existing latently infected cells.

Supplementing allo-HSCTs with cellular therapies may improve viral reservoir clearance [42, 43]. Donor lymphocyte infusions (DLIs) can drive relapsing hematologic tumors into remission [44]. Conceivably, incorporating DLIs into allo-HSCT treatment plans could similarly augment GvVR responses [25]. However, since DLIs involve transferring polyclonal T cells, they also carry a significant risk of GvHD [45]. Therefore, targeted cellular immunotherapies maximizing GvVR effects while minimizing GvHD side effects are preferred. For instance, repurposing cancer adoptive cell therapies may boost GvVR activity [46–48]. HIV-specific chimeric antigen receptor (CAR) T- and NK-cell therapies are promising HIV cure strategies [49] despite early iterations having limited impact on HIV reservoirs [50–53]. Alternatively, focusing T cells on minor histocompatibility antigens (mHAgs) exclusively expressed by host leukocytes may promote GvVR effects without causing GvHD [54]. We recently identified a mHAg in APOBEC3C that is expressed preferentially by MCM immune cells [55]. Hence, this epitope may serve as a model antigen for testing mHAg-targeted cellular immunotherapies in NHPs and examining their capacity to eradicate viral reservoirs.

In this study, we used the pathogenic SIV-HIV chimera SHIV162P3 to model HIV infections. This recombinant virus contains the *env* gene from the clade B isolate HIV-1 SF162 inserted into the SIVmac239 genomic backbone [56]. Importantly, SHIV162P3 exclusively uses CCR5 as a coreceptor, similar to most transmitted HIV-1 strains [35, 57]. This coreceptor usage contrasts with common SIV strains, which can use alternate chemokine receptors and G-protein coupled receptors for cell entry [58, 59]. However, SHIV162P3 replication is less robust than most SIV strains, resulting in lower chronic phase viral loads [60, 61]. Indeed, MCM-A and -B spontaneously suppressed viremia to approximately 10^3^ vRNA copies/ml plasma by four months post-infection. Nevertheless, SHIV162P3 established pre-transplant viral reservoirs in our MCMs consistent with previous SHIV/ SIV studies [36, 40, 62, 63].

Our study has several limitations. First, its small size and lack of a control group transplanted with TCR α/β-replete HSCs make it difficult to draw firm conclusions from the results. Second, ART was started 3-6 weeks before transplantation. While this provided sufficient time to suppress SHIV replication, it does not reflect the prolonged ART typical of HIV-infected allo-HSCT recipients [4, 5, 8, 14, 15]. Third, we could not determine whether virally infected cells were of the donor or recipient origin post-transplant. A study by Colonna *et al.* exploited haploidentical allo-HSCTs in rhesus macaques to sort donor and recipient cells based on differential MHC expression, finding that most SHIV-infected cells were from the recipients early after transplantation [40]. However, our MHC-matched MCMs lacked unique surface markers to distinguish donor and recipient cells, prohibiting similar analyses. Lastly, we achieved poor donor cell engraftment, hindering post-transplant analyses and potentially limiting SHIV reservoir depletion. Consequently, we could not quantify donor chimerism in tissues or measure post-transplant antiviral and GvVR immune responses. The limited lymphocyte recovery also hampered our ability to longitudinally measure SHIV-infected cells post-transplant. Therefore, less toxic pre-transplant conditioning regimens, higher HSC doses, or T-cell replete HSCs may be necessary for sustaining donor cell engraftment. To this end, Burwitz *et al.* established an allo-HSCT model in MHC-matched MCMs that achieves durable post-transplant donor chimerism with reduced-intensity conditioning but has an increased risk of GvHD [41]. Thus, allo-HSCTs in NHPs will likely need to balance full donor engraftment with suppressing GvHD to dissect the mechanisms of GvVR effects.

## Conclusions

In summary, MHC-matched bone marrow transplants in ART-suppressed SHIV-infected MCMs reduced, but did not eliminate, viral reservoirs early after transplantation. However, depleting TCRα/β T cells from the HSCs and post-transplant prophylaxis prevented high-grade GvHD. Our results suggest that GvVR responses and sustained ART after allo-HSCTs may be needed to significantly deplete viral reservoirs.

## Methods

### Ethics statement and animal care

The staff at the Wisconsin National Primate Research Center (WNPRC) cared for the Cynomolgus macaques (*Macaca fascicularis*) according to the regulations and guidelines of the University of Wisconsin Institutional Animal Care and Use Committee, which approved this study (protocol g005424) following the recommendations of the Weatherall Report and the principles described in the National Research Council’s Guide for the Care and Use of Laboratory Animals. The MCMs were closely monitored for signs of stress or pain. In consultation with the WNPRC veterinarians, the MCMs were euthanized if they developed an untreatable opportunistic infection, inappetence, and/or progressive decline in condition.

### MHC typing

MHC genotyping was performed by Genetic Services at WNPRC as previously described [64]. Briefly, genomic DNA (gDNA) was isolated from whole blood or PBMCs and used as templates in polymerase chain reactions (PCRs) with primers flanking exon 2 of MHC class I (Mafa-A, Mafa-B, Mafi-I, and Mafa-E) and class II (Mafa-DRB, Mafa-DQA, Mafa-DQB, Mafa-DPA, and Mafa-DPB) loci. The PCR reactions were generated with a Fluidigm Access Array, which permitted all reactions to be multiplexed in a single experiment. After cleanup with AMPure beads (Beckman Coulter), the amplicons were sequenced on an Illumina MiSeq. The resulting sequences were mapped against a custom database of MCM class I and II sequences to assign one of seven MCM haplotypes (M1-M7).

### SHIV infections and viral loads

We inoculated the MCMs intravenously with 500 median tissue culture infectious doses (TCID_50_) of SHIV162P3 [35]. Plasma viral loads were measured via quantitative real-time polymerase chain reactions (qRT-PCRs) as previously described [65, 66]. Briefly, vRNA was isolated from plasma using the Maxwell Viral Total Nucleic Acid Purification kit (Promega) on a Maxwell 48 RSC instrument (Promega). Next, vRNA was reverse transcribed into complementary DNA (cDNA), amplified using the TaqMan Fast Virus 1-Step Master Mix qRT-PCR kit (Invitrogen) on a LightCycler 480 or LC96 instrument (Roche), and quantified by interpolation onto a standard curve of 10-fold serial dilutions of an SIV *gag in vitro* transcript. The assay limit of detection was 100 vRNA copies/ml.

### Antiretroviral treatment

We treated the SHIV-infected MCMs with a combination of reverse transcriptase inhibitors tenofovir (PMPA) and emtricitabine (FTC) and integrase inhibitor raltegravir (RAL). PMPA and FTC were pre-formulated at concentrations of 20 mg/ml and 40 mg/ml in water containing sodium hydroxide (NaOH) and administered subcutaneously once daily at 1 ml/kg of body weight. 100 mg of RAL was mixed twice daily with food [67]. Gilead Sciences kindly provided PMPA and FTC, and Merck kindly provided RAL through material transfer agreements.

### Donor bone marrow collection and TCRα/β cell depletion

We prepared the HSCs as previously described [34]. Briefly, we collected BM from sex and MHC-matched MCMs (Table 1) by aspirating up to 5 mL from four sites, 20 mL total, and removed red blood cells using ACK lysis buffer (Thermo Fisher Scientific). Next, we washed the cells twice with phosphate-buffered saline and incubated them with anti-TCRα/β allophycocyanin (APC) antibodies (clone R73, BioLegend) for 20 minutes, followed by incubating with anti-APC microbeads (Miltenyi Biotec) for an additional 20 minutes at 4°C.

Finally, we passed the stained cells through two LS columns (Miltenyi Biotec) stacked on top of one another, collecting both negative and positive cell fractions, and cryopreserving them at 30 million/mL in serum-free expansion medium (SFEM; Stem Cell Technologies) containing 5% fetal bovine serum and 10% dimethylsulfoxide (DMSO) until further use.

### Bone marrow transplants and care

First, we treated the MCMs with daily oral antibiotics trimethoprim (200 mg) and sulfamethoxazole (40 mg) to decontaminate the gut, starting seven days pre-transplant (−7) and continuing until day −1. Next, we started the MCMs on systemic bacterial prophylaxis by administering ceftriaxone (50 mg/kg) on day −2; MCM-D was switched to cefazolin (25 mg/kg twice daily) at day −1, and maintaining treatment until absolute neutrophil counts stabilized. On day −2, the MCMs received five separate 1 Gy fractions of TBI on two consecutive days (10 Gy total), sparing the lungs and eyes as previously described [34]. One day before transplantation, we thawed cryopreserved TCRα/β-depleted BM fractions at 37°C and cultured them overnight in SFEM containing 100 ng/mL human stem cell factor (SCF, Peprotech), 100 ng/mL human Fms-related tyrosine kinase 3 (FLT3) ligand (Peprotech), and 50 ng/mL human thrombopoietin (TPO, Peprotech). On the day of transplantation, we resuspended the BM cells in 15 mL of Plasma-Lyte (Baxter), a nonpyrogenic isotonic solution, supplemented with 2% autologous serum and 5 U/mL heparin, and infused the cell suspension intravenously. Next, we prophylactically treated the MCMs for GvHD using cyclophosphamide (50 mg/kg) on days 4 and 5 post-transplant, followed by twice-daily tacrolimus (0.01 mg/kg) starting at day 5, maintaining tacrolimus serum levels between 5-15 ng/mL. To prevent fungal infections, we treated the MCMs with once-daily fluconazole (5 mg/kg) starting on day 0. Lastly, we treated MCMs C and D with oral eltrombopag (1.5 mg/kg) and N-acetyl L-cysteine (50 mg/kg), respectively, to support platelet engraftment. See Supplementary Table 1 for treatment details.

### Measuring peripheral leukocyte populations and GvHD

Following transplantation, we collected peripheral blood from the MCMs twice per week for the first month, followed by bi-weekly blood draws for the next two months and monthly thereafter. We used an XS-1000i automated hematology analyzer (Sysmex) to measure platelets and leukocyte populations. In addition, GvHD was evaluated as previously described [68].

We quantified CD4^+^ and CD8^+^ T cells by incubating 100 μl of whole blood with anti-CD8-Brilliant Violet 421 (clone SK1; BioLegend), anti-CD4-APC (clone L200; BD-Pharmingen), and anti-CD3-Alexa 700 (clone SP34; BD-Pharmingen) antibodies with Live/Dead Fixable Near-IR vital dye (ThermoFisher Scientific) for 30 min at 4°C. Next, we lysed the red blood cells by adding 1 ml of FACS Lysing Solution (BD Biosciences) and incubated the samples for 10 minutes at room temperature. We washed the cells three times with FACS buffer (phosphate-buffered saline supplemented with 2% fetal calf serum), fixed with 2% paraformaldehyde, and ran the samples on a BD-LSR-II flow cytometer using FACSDiva software (Becton Dickinson). Finally, we analyzed the flow cytometry data using FlowJo software (Treestar), determining the absolute CD4^+^ and CD8^+^ T cell counts by multiplying the frequency of CD4^+^ or CD8^+^ T cells (singlets, live, CD3^+^, CD4^+^ or CD8^+^) by the white blood counts per microliter of blood from the matching complete blood counts (Supplementary Figure 1).

### Lymphocyte isolation from tissues

We isolated lymphocytes from the blood, lymph nodes, spleen, and bone marrow as previously described [69]. Briefly, we isolated PBMCs from EDTA-anticoagulated blood by density gradient centrifugation using Ficoll-Paque PLUS (GE Healthcare), removing contaminating red blood cells with ACK lysis buffer. We collected peripheral lymph nodes before ART (MCM A and B: 77 days post-infection (dpi); MCM-C: unavailable; and MCM-D: 34 dpi), after ART but before allo-HSCT (MCM-A: 23 days post-ART; MCM-B: 24 days post-ART; MCM-C: 18 days post-ART; and MCM-D: 12 days post-ART). Additionally, we collected lymph nodes from MCM-A 62 days post-transplant but before ART cessation and the necropsies for MCM C and D, which occurred on days 46 and 39 post-transplant, respectively, while they were on ART. We were unable to collect lymph nodes at MCM-B’s necropsy for analysis.

We isolated lymphocytes from the lymph nodes, spleen, and bone marrow by dicing the tissues with scalpels and forcing the cells through 100-μm cell strainers to remove connective tissues. Lymphocytes were enriched from the cell suspensions using Ficoll-Paque PLUS density gradient centrifugation.

### Measuring donor chimerism

We identified SNPs that distinguished the donor and recipient MCMs using a panel of 12 SNPs (see Supplementary Table 2) and the rhAMP SNP Genotyping System (Integrated DNA Technologies) as described previously [34, 70], giving preference to homozygous/homozygous mismatches. We performed the rhAMP assays in triplicate using a LightCycler 96 Instrument (Roche Molecular Systems) in the endpoint analysis mode.

We quantified donor chimerism in whole PBMCs or immune subsets by sequencing the diagnostic SNPs with an Illumina MiSeq using previously described primer sets, PCR conditions, and analysis pipeline (see also Supplementary Table 3 and Supplementary Table 4) [34]. To isolate specific immune subsets, we stained PBMCs with anti-CD3 FITC (clone SP34; BD Pharmingen), anti-CD14 PerCP-Cy5.5 (clone M5E2; BioLegend), anti-CD20 APC (clone 2H7; BioLegend), anti-CD45 PE (clone D058-1283; BD Pharmingen), and Live/Dead Fixable Near-IR vital dye (ThermoFisher Scientific) and sorted into monocyte (singlets, live, lymphocytes, CD45^mid^, CD14^+^ cells), T lymphocyte (singlets, live, lymphocytes, CD14^-^, CD45^hi^, CD3^+^), or B lymphocyte (singlets, live, lymphocytes, CD14^-^, CD45^hi^, CD20^+^) populations using a BD FACSJazz cell sorter. We used between 179 and 61,251 cells for gDNA extraction and sequencing for each sorted population.

### Quantifying cell-associated viral DNA

We extracted gDNA from tissues or PBMCs using the DNeasy Blood & Tissue Kit (Qiagen) per the manufacturer’s instructions, except eluting the DNA twice with separate 75 μL of molecular grade water (150 μl total volume). We determined the concentration of DNA using a Nanodrop spectrophotometer (ThermoFisher Scientific), and we digested up to 3 μg of DNA with 3 μl of EcoRI-HF restriction enzyme (New England Biolabs) per 50 μl.

We performed two digital droplet PCR (ddPCR) reactions for each sample: one quantifying vDNA and the other determining total cell number. First, we quantified vDNA using the primers and probes for SIV *gag* and the cycling conditions described by Gama *et al.* [71]. Second, we normalized vDNA copies to cell numbers by quantifying the RNase P (RPP30) gene, as previously described [72]. We generated droplets and read the samples on a QX200 ddPCR system (Bio-Rad). We quantified the target genes in duplicate for each sample across at least two independent runs.

## Abbreviations

Allo-HSCTs: Allogeneic hematopoietic stem cell transplants
ART: Antiretroviral therapy
HIV: Human immunodeficiency virus
SHIV: Simian-human immunodeficiency virus
MCMs: Mauritian cynomolgus macaques
GvH: Graft-versus-host
GvHD: Graft-versus-host disease
GvVR: Graft-versus-viral reservoir
CCR5: C-C chemokine receptor type 5
CXCR4: C-X-C chemokine receptor type 4
CCR5Δ32: C-C chemokine receptor type 5 with 32 base-pair deletion
MHC: Major histocompatibility complex
NHPs: Nonhuman primates
BM: Bone marrow
p.i.: Post-infection
Gy: Gray
WBCs: White blood cells
ATIs: Analytical treatment interruptions
DLIs: Donor lymphocyte infusions
CAR: Chimeric antigen receptor
mHAgs: Minor histocompatibility antigens
WNPRC: Wisconsin national primate research center
gDNA: Genomic deoxyribonucleic acid
PCRs: Polymerase chain reactions
Mafa: Macaca fascicularis
TCID_50_: Median tissue culture infectious doses
qRT-PCRs: Quantitative real-time polymerase chain reactions
cDNA: Complementary deoxyribonucleic acid
PMPA: Tenofovir (9-[9(R)-2-(phosphonomethoxy)propyl]adenine
FTC: Emtricitabine
RAL: Raltegravir
SFEM: Serum-free expansion medium
TBI: Total body irradiation
SCF: Stem cell factor
FLT3: Fms-related tyrosine kinase 3
TPO: Thrombopoietin
PBMCs: Peripheral blood mononuclear cells
SNPs: Single-nucleotide polymorphisms
vDNA: viral deoxyribonucleic acids
ddPCR: Digital droplet polymerase chain reaction
RPP30: RNase P subunit p30

## Authors’ contributions

M.R.R., T. G., and I.I.S.: Conceptualization and study design; J.T.W and S.S.D.: coordinated the animal experiments; J.T.W.: processed samples and performed flow cytometry; S.S.D., K.S., and A.K.: prepared TCR α/β-depleted HSCs; L.M.M.: performed SHIV DNA measurements; S.B. and L.E.K.: performed chimerism analysis; J.C.: contributed to treatment design and handled animals; C.M.C. and P.H.: advised on the allo-HSCT platform, GvHD prophylaxis, and critically reviewed the manuscript; M.R.R.: wrote the manuscript. All authors revised the manuscript and approved its submission.

## Acknowledgments

We thank the WNPRC veterinary staff for their assistance.

## References

1. Dickinson AM, Norden J, Li S et al. Graft-versus-Leukemia Effect Following Hematopoietic Stem Cell Transplantation for Leukemia. Front Immunol. 2017;8:496.

2. Falkenburg JH, Warren EH. Graft versus leukemia reactivity after allogeneic stem cell transplantation. Biol Blood Marrow Transplant. 2011;17:S33–S38.

3. Warren EH, Fujii N, Akatsuka Y et al. Therapy of relapsed leukemia after allogeneic hematopoietic cell transplantation with T cells specific for minor histocompatibility antigens. Blood. 2010;115:3869–3878.

4. Cummins NW, Rizza S, Litzow MR et al. Extensive virologic and immunologic characterization in an HIV-infected individual following allogeneic stem cell transplant and analytic cessation of antiretroviral therapy: A case study. PLoS Med. 2017;14:e1002461.

5. Gupta RK, Abdul-Jawad S, McCoy LE et al. HIV-1 remission following CCR5Δ32/Δ32 haematopoietic stem-cell transplantation. Nature. 2019

6. Gupta RK, Peppa D, Hill AL et al. Evidence for HIV-1 cure after CCR5Δ32/Δ32 allogeneic haemopoietic stem-cell transplantation 30 months post analytical treatment interruption: a case report. Lancet HIV. 2020

7. Henrich TJ, Hu Z, Li JZ et al. Long-term reduction in peripheral blood HIV type 1 reservoirs following reduced-intensity conditioning allogeneic stem cell transplantation. J Infect Dis. 2013;207:1694–1702.

8. Hutter G, Nowak D, Mossner M et al. Long-term control of HIV by CCR5 Delta32/Delta32 stem-cell transplantation. N Engl J Med. 2009;360:692–698.

9. Koelsch KK, Rasmussen TA, Hey-Nguyen WJ et al. Impact of Allogeneic Hematopoietic Stem Cell Transplantation on the HIV Reservoir and Immune Response in 3 HIV-Infected Individuals. J Acquir Immune Defic Syndr. 2017;75:328–337.

10. Prator CA, Donatelli J, Henrich TJ. From Berlin to London: HIV-1 Reservoir Reduction Following Stem Cell Transplantation. Curr HIV/AIDS Rep. 2020;17:385–393.

11. Smiley ST, Singh A, Read SW et al. Progress toward curing HIV infections with hematopoietic stem cell transplantation. Clin Infect Dis. 2015;60:292–297.

12. Hutter G. Stem cell transplantation in strategies for curing HIV/AIDS. AIDS Res Ther. 2016;13:31.

13. Allers K, Hutter G, Hofmann J et al. Evidence for the cure of HIV infection by CCR5Delta32/Delta32 stem cell transplantation. Blood. 2011;117:2791–2799.

14. Salgado M, Kwon M, Gálvez C et al. Mechanisms that contribute to a profound reduction of the HIV-1 reservoir after allogeneic stem cell transplant. Ann Intern Med. 2018;169:674–683.

15. Henrich TJ, Hanhauser E, Marty FM et al. Antiretroviral-free HIV-1 remission and viral rebound after allogeneic stem cell transplantation: report of 2 cases. Ann Intern Med. 2014;161:319–327.

16. Hill AL, Rosenbloom DI, Goldstein E et al. Real-Time Predictions of Reservoir Size and Rebound Time during Antiretroviral Therapy Interruption Trials for HIV. PLoS Pathog. 2016;12:e1005535.

17. Eberhard JM, Angin M, Passaes C et al. Vulnerability to reservoir reseeding due to high immune activation after allogeneic hematopoietic stem cell transplantation in individuals with HIV-1. Sci Transl Med. 2020;12:eaay9355.

18. Gyurkocza B, Sandmaier BM. Conditioning regimens for hematopoietic cell transplantation: one size does not fit all. Blood. 2014;124:344–353.

19. Cillo AR, Krishnan S, McMahon DK, Mitsuyasu RT, Para MF, Mellors JW. Impact of chemotherapy for HIV-1 related lymphoma on residual viremia and cellular HIV-1 DNA in patients on suppressive antiretroviral therapy. PLoS One. 2014;9:e92118.

20. Cillo AR, Krishnan A, Mitsuyasu RT et al. Plasma viremia and cellular HIV-1 DNA persist despite autologous hematopoietic stem cell transplantation for HIV-related lymphoma. J Acquir Immune Defic Syndr. 2013;63:438–441.

21. Gabarre J, Marcelin AG, Azar N et al. High-dose therapy plus autologous hematopoietic stem cell transplantation for human immunodeficiency virus (HIV)-related lymphoma: results and impact on HIV disease. Haematologica. 2004;89:1100–1108.

22. Henrich TJ, Hobbs KS, Hanhauser E et al. Human Immunodeficiency Virus Type 1 Persistence Following Systemic Chemotherapy for Malignancy. J Infect Dis. 2017;216:254–262.

23. Mavigner M, Watkins B, Lawson B et al. Persistence of virus reservoirs in ART-treated SHIV-infected rhesus macaques after autologous hematopoietic stem cell transplant. PLoS Pathog. 2014;10:e1004406.

24. Simonelli C, Zanussi S, Pratesi C et al. Immune recovery after autologous stem cell transplantation is not different for HIV-infected versus HIV-uninfected patients with relapsed or refractory lymphoma. Clin Infect Dis. 2010;50:1672–1679.

25. Serrano D, Miralles P, Balsalobre P et al. Graft-versus-tumor effect after allogeneic stem cell transplantation in HIV-positive patients with high-risk hematologic malignancies. AIDS Res Hum Retroviruses. 2013;29:1340–1345.

26. Griffioen M, van Bergen CA, Falkenburg JH. Autosomal Minor Histocompatibility Antigens: How Genetic Variants Create Diversity in Immune Targets. Front Immunol. 2016;7:100.

27. Warren EH, Deeg HJ. Dissecting graft-versus-leukemia from graft-versus-host-disease using novel strategies. Tissue Antigens. 2013;81:183–193.

28. Lawler SH, Sussman RW, Taylor LL. Mitochondrial DNA of the Mauritian macaques (*Macaca fascicularis*): an example of the founder effect. Am J Phys Anthropol. 1995;96:133–141.

29. Budde ML, Wiseman RW, Karl JA, Hanczaruk B, Simen BB, O’Connor DH. Characterization of Mauritian cynomolgus macaque major histocompatibility complex class I haplotypes by high-resolution pyrosequencing. Immunogenetics. 2010;62:773–780.

30. O’Connor SL, Blasky AJ, Pendley CJ et al. Comprehensive characterization of MHC class II haplotypes in Mauritian cynomolgus macaques. Immunogenetics. 2007;59:449–462.

31. Wiseman RW, Wojcechowskyj JA, Greene JM et al. Simian immunodeficiency virus SIVmac239 infection of major histocompatibility complex-identical cynomolgus macaques from Mauritius. J Virol. 2007;81:349–361.

32. Antony JM, MacDonald KS. A critical analysis of the cynomolgus macaque, Macaca fascicularis, as a model to test HIV-1/SIV vaccine efficacy. Vaccine. 2015;33:3073–3083.

33. Burwitz BJ, Pendley CJ, Greene JM et al. Mauritian cynomolgus macaques share two exceptionally common major histocompatibility complex class I alleles that restrict simian immunodeficiency virus-specific CD8+ T cells. J Virol. 2009;83:6011–6019.

34. D’Souza SS, Bennett S, Kumar A et al. Transplantation of T-cell receptor α/β-depleted allogeneic bone marrow in nonhuman primates. Exp Hematol. 2021;93:44–51.

35. Harouse JM, Gettie A, Eshetu T et al. Mucosal transmission and induction of simian AIDS by CCR5-specific simian/human immunodeficiency virus SHIV(SF162P3). J Virol. 2001;75:1990–1995.

36. Cadena AM, Ventura JD, Abbink P et al. Persistence of viral RNA in lymph nodes in ART-suppressed SIV/SHIV-infected Rhesus Macaques. Nat Commun. 2021;12:1474.

37. Wong JK, Yukl SA. Tissue reservoirs of HIV. Curr Opin HIV AIDS. 2016;11:362–370.

38. Julg B, Dee L, Ananworanich J et al. Recommendations for analytical antiretroviral treatment interruptions in HIV research trials-report of a consensus meeting. Lancet HIV. 2019;6:e259–e268.

39. Lau JSY, Smith MZ, Lewin SR, McMahon JH. Clinical trials of antiretroviral treatment interruption in HIV-infected individuals. AIDS. 2019;33:773–791.

40. Colonna L, Peterson CW, Schell JB et al. Evidence for persistence of the SHIV reservoir early after MHC haploidentical hematopoietic stem cell transplantation. Nat Commun. 2018;9:4438.

41. Burwitz BJ, Wu HL, Abdulhaqq S et al. Allogeneic stem cell transplantation in fully MHC-matched Mauritian cynomolgus macaques recapitulates diverse human clinical outcomes. Nat Commun. 2017;8:1418.

42. Ward AR, Mota TM, Jones RB. Immunological approaches to HIV cure. Semin Immunol. 2020101412.

43. Zhou Y, Maldini CR, Jadlowsky J, Riley JL. Challenges and Opportunities of Using Adoptive T-Cell Therapy as Part of an HIV Cure Strategy. J Infect Dis. 2021;223:38–45.

44. Kolb HJ, Schattenberg A, Goldman JM et al. Graft-versus-leukemia effect of donor lymphocyte transfusions in marrow grafted patients. Blood. 1995;86:2041–2050.

45. Frey NV, Porter DL. Graft-versus-host disease after donor leukocyte infusions: presentation and management. Best Pract Res Clin Haematol. 2008;21:205–222.

46. Mylvaganam G, Yanez AG, Maus M, Walker BD. Toward T Cell-Mediated Control or Elimination of HIV Reservoirs: Lessons From Cancer Immunology. Front Immunol. 2019;10:2109.

47. Rust BJ, Kiem HP, Uldrick TS. CAR T-cell therapy for cancer and HIV through novel approaches to HIV-associated haematological malignancies. Lancet Haematol. 2020;7:e690–e696.

48. Yang H, Wallace Z, Dorrell L. Therapeutic Targeting of HIV Reservoirs: How to Give T Cells a New Direction. Front Immunol. 2018;9:2861.

49. Mu W, Carrillo MA, Kitchen SG. Engineering CAR T Cells to Target the HIV Reservoir. Front Cell Infect Microbiol. 2020;10:410.

50. Deeks SG, Wagner B, Anton PA et al. A phase II randomized study of HIV-specific T-cell gene therapy in subjects with undetectable plasma viremia on combination antiretroviral therapy. Mol Ther. 2002;5:788–797.

51. Mitsuyasu RT, Anton PA, Deeks SG et al. Prolonged survival and tissue trafficking following adoptive transfer of CD4zeta gene-modified autologous CD4(+) and CD8(+) T cells in human immunodeficiency virus-infected subjects. Blood. 2000;96:785–793.

52. Scholler J, Brady TL, Binder-Scholl G et al. Decade-long safety and function of retroviral-modified chimeric antigen receptor T cells. Sci Transl Med. 2012;4:132ra53.

53. Walker RE, Bechtel CM, Natarajan V et al. Long-term in vivo survival of receptor-modified syngeneic T cells in patients with human immunodeficiency virus infection. Blood. 2000;96:467–474.

54. Summers C, Sheth V, Bleakley M. Minor histocompatibility antigen-specific T cells. Frontiers in Pediatrics. 2020

55. Weinfurter JT, Graham ME, Ericsen AJ et al. Identifying a Minor Histocompatibility Antigen in Mauritian Cynomolgus Macaques Encoded by APOBEC3C. Front Immunol. 2020;11:586251.

56. Luciw PA, Pratt-Lowe E, Shaw KE, Levy JA, Cheng-Mayer C. Persistent infection of rhesus macaques with T-cell-line-tropic and macrophage-tropic clones of simian/human immunodeficiency viruses (SHIV). Proc Natl Acad Sci U S A. 1995;92:7490–7494.

57. Moore JP, Kitchen SG, Pugach P, Zack JA. The CCR5 and CXCR4 coreceptors--central to understanding the transmission and pathogenesis of human immunodeficiency virus type 1 infection. AIDS Res Hum Retroviruses. 2004;20:111–126.

58. Sina ST, Ren W, Cheng-Mayer C. Coreceptor use in nonhuman primate models of HIV infection. J Transl Med. 2011;9 Suppl 1:S7.

59. Zhang Y, Lou B, Lal RB, Gettie A, Marx PA, Moore JP. Use of inhibitors to evaluate coreceptor usage by simian and simian/human immunodeficiency viruses and human immunodeficiency virus type 2 in primary cells. J Virol. 2000;74:6893–6910.

60. Pahar B, Lackner AA, Piatak M et al. Control of viremia and maintenance of intestinal CD4(+) memory T cells in SHIV(162P3) infected macaques after pathogenic SIV(MAC251) challenge. Virology. 2009;387:273–284.

61. Xu H, Wang X, Morici LA, Pahar B, Veazey RS. Early divergent host responses in SHIVsf162P3 and SIVmac251 infected macaques correlate with control of viremia. PLoS One. 2011;6:e17965.

62. Siddiqui S, Perez S, Gao Y et al. Persistent Viral Reservoirs in Lymphoid Tissues in SIV-Infected Rhesus Macaques of Chinese-Origin on Suppressive Antiretroviral Therapy. Viruses. 2019;11:E105.

63. Abreu CM, Veenhuis RT, Avalos CR et al. Infectious Virus Persists in CD4^+^ T Cells and Macrophages in Antiretroviral Therapy-Suppressed Simian Immunodeficiency Virus-Infected Macaques. J Virol. 2019;93:e00065–19.

64. Wiseman RW, Karl JA, Bohn PS, Nimityongskul FA, Starrett GJ, O’Connor DH. Haplessly hoping: macaque major histocompatibility complex made easy. ILAR J. 2013;54:196–210.

65. Cline AN, Bess JW, Piatak M, Jr., Lifson JD. Highly sensitive SIV plasma viral load assay: practical considerations, realistic performance expectations, and application to reverse engineering of vaccines for AIDS. J Med Primatol. 2005;34:303–312.

66. Friedrich TC, Valentine LE, Yant LJ et al. Subdominant CD8+ T-cell responses are involved in durable control of AIDS virus replication. J Virol. 2007;81:3465–3476.

67. Lewis MG, Norelli S, Collins M et al. Response of a simian immunodeficiency virus (SIVmac251) to raltegravir: a basis for a new treatment for simian AIDS and an animal model for studying lentiviral persistence during antiretroviral therapy. Retrovirology. 2010;7:21.

68. Miller WP, Srinivasan S, Panoskaltsis-Mortari A et al. GVHD after haploidentical transplantation: a novel, MHC-defined rhesus macaque model identifies CD28-CD8+ T cells as a reservoir of breakthrough T-cell proliferation during costimulation blockade and sirolimus-based immunosuppression. Blood. 2010;116:5403–5418.

69. Reynolds MR, Rakasz E, Skinner PJ et al. CD8+ T-lymphocyte response to major immunodominant epitopes after vaginal exposure to simian immunodeficiency virus: too late and too little. J Virol. 2005;79:9228–9235.

70. Maufort JP, Israel JS, Brown ME et al. Major Histocompatibility Complex-Matched Arteries Have Similar Patency to Autologous Arteries in a Mauritian Cynomolgus Macaque Major Histocompatibility Complex-Defined Transplant Model. J Am Heart Assoc. 2019;8:e012135.

71. Gama L, Abreu CM, Shirk EN et al. Reactivation of simian immunodeficiency virus reservoirs in the brain of virally suppressed macaques. AIDS. 2017;31:5–14.

72. Dyavar SR, Ye Z, Byrareddy SN et al. Normalization of cell associated antiretroviral drug concentrations with a novel RPP30 droplet digital PCR assay. Sci Rep. 2018;8:3626.

