## Supplementary Figure 1 for "MHC-matched allogeneic bone marrow transplants fail to eliminate SHIV-infected cells from ART-suppressed Mauritian cynomolgus macaques"

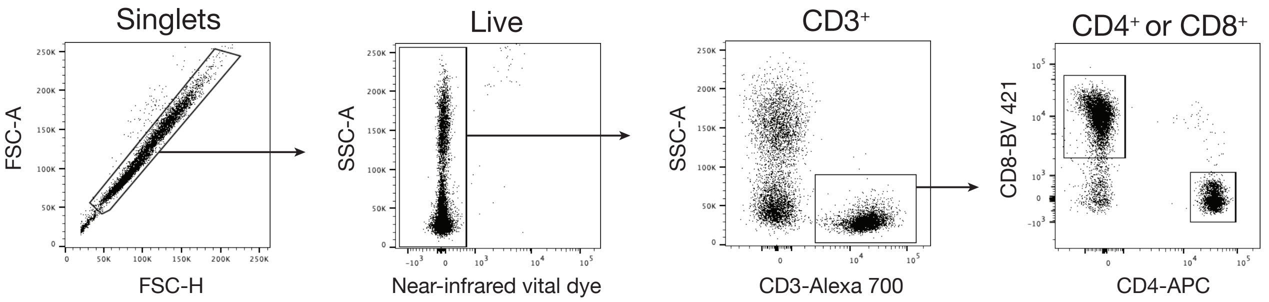


**Supplementary figure 1.** Flow cytometric gating strategy for determining the frequency of $CD4^+$ and $CD8^+$ T cells in the blood, progressively selecting singlets, live cells, $CD3^+$, and either $CD4^+$ or $CD8^+$ cells. Representative data from a pre-transplant sample.
