## Supplementary Table 1 for "MHC-matched allogeneic bone marrow transplants fail to eliminate SHIV-infected cells from ART-suppressed Mauritian cynomolgus macaques"

**Supplementary Table 1.** Transplant recipients' treatment regimens.

MCM-A

days post-transplant

| Treatment | Purpose | Dose | Route | -30 | .. | - 7 | .. | -2 | -1 | 0 | 1 | 2 | 3 | 4 | 5 | .. | 9 | .. | 19 | .. | 26 | .. | 42 | .. | 58 | .. | 62 | 63 | .. | 74 |
|---|---|---|---|---|---|---|---|---|---|---|---|---|---|---|---|---|---|---|---|---|---|---|---|---|---|---|---|---|---|---|
| ART- TDF<br>FTC | Anti-<br>retroviral | 20mg/kg<br>40mg/kg | SQ<br>SQ | | | | | | | | | | | | | | | | | | | | | | | | | | | |
| ART- RAL | Anti-<br>retroviral | 100mg | oral | | | | | | | | | | | | | | | | | | | | | | | | | | | |
| Trimethoprim/<br>sulfamethoxazole | Gut<br>decontamination | 200mg<br>40mg | oral | | | | | | | | | | | | | | | | | | | | | | | | | | | |
| Ceftriaxone | Antibiotic | 50<br>mg/kg | IM | | | | | | | | | | | | | | | | | | | | | | | | | | | |
| Total body irradiation | Immune<br>conditioning | 5Gy for 2<br>days | - | | | | | | | | | | | | | | | | | | | | | | | | | | | |
| TCRα/β-depleted<br>HSCs | | 2x10^8<br>cells | IV | | | | | | | | | | | | | | | | | | | | | | | | | | | |
| Cyclophosphamide | Immune<br>suppressant | 50<br>mg/kg | IV | | | | | | | | | | | | | | | | | | | | | | | | | | | |
| Tacrolimus | Immune<br>suppressant | 0.01<br>mg/kg | IM | | | | | | | | | | | | | | | | | | | | | | | | | | | |
| Fluconazole | Anti-fungal | 5<br>mg/kg | oral | | | | | | | | | | | | | | | | | | | | | | | | | | | |
| Blood transfusion | | | IV | | | | | | | | | | | | | | | | | | | | | | | | | | | |
| Necropsy | | | | | | | | | | | | | | | | | | | | | | | | | | | | | | |

MCM-B

days post-transplant

| Treatment | Purpose | Dose | Route | -42 | .. | - 7 | .. | -2 | -1 | 0 | 1 | 2 | 3 | 4 | 5 | .. | 12 | .. | 19 | .. | 26 | .. | 33 | .. | 38 |
|---|---|---|---|---|---|---|---|---|---|---|---|---|---|---|---|---|---|---|---|---|---|---|---|---|---|
| ART- TDF<br>FTC | Anti-<br>retroviral | 20mg/kg<br>40mg/kg | SQ<br>SQ | | | | | | | | | | | | | | | | | | | | | | |
| ART- RAL | Anti-<br>retroviral | 100mg | oral | | | | | | | | | | | | | | | | | | | | | | |
| Trimethoprim/<br>sulfamethoxazole | Gut<br>decontamination | 200mg<br>40mg | oral | | | | | | | | | | | | | | | | | | | | | | |
| Ceftriaxone | Antibiotic | 50<br>mg/kg | IM | | | | | | | | | | | | | | | | | | | | | | |
| Total body irradiation | Immune<br>conditioning | 5Gy for 2<br>days | - | | | | | | | | | | | | | | | | | | | | | | |
| TCRα/β-depleted<br>cells | | 8.5x10^8<br>cells | IV | | | | | | | | | | | | | | | | | | | | | | |
| Cyclophosphamide | Immune<br>suppressant | 50<br>mg/kg | IV | | | | | | | | | | | | | | | | | | | | | | |
| Tacrolimus | Immune<br>suppressant | 0.01<br>mg/kg | IM | | | | | | | | | | | | | | | | | | | | | | |
| Fluconazole | Anti-<br>fungal | 5<br>mg/kg | oral | | | | | | | | | | | | | | | | | | | | | | |
| Blood transfusion | | | IV | | | | | | | | | | | | | | | | | | | | | | |
| Necropsy | | | | | | | | | | | | | | | | | | | | | | | | | |

### MCM-C

days post-transplant

| Treatment | Purpose | Dose | Route | -24 | .. | - 7 | .. | -2 | -1 | 0 | 1 | 2 | 3 | 4 | 5 | .. | 11 | .. | 19 | .. | 28 | .. | 34 | 35 | .. | 37 | .. | 42 | .. | 45 |
|---|---|---|---|---|---|---|---|---|---|---|---|---|---|---|---|---|---|---|---|---|---|---|---|---|---|---|---|---|---|---|
| ART- TDF FTC | Anti-retroviral | 20mg/kg 40mg/kg | SQ SQ | | | | | | | | | | | | | | | | | | | | | | | | | | | |
| ART-RAL | Anti-retroviral | 100mg | oral | | | | | | | | | | | | | | | | | | | | | | | | | | | |
| Trimethoprim/ sulfamethoxazole | Gut decontamination | 200mg 40mg | oral | | | | | | | | | | | | | | | | | | | | | | | | | | | |
| Ceftriaxone | Antibiotic | 50 mg/kg | IM | | | | | | | | | | | | | | | | | | | | | | | | | | | |
| Total body irradiation | Immune conditioning | 5Gy for 2 days | - | | | | | | | | | | | | | | | | | | | | | | | | | | | |
| TCRα/β-depleted cells | | 7.1x10^8 cells | IV | | | | | | | | | | | | | | | | | | | | | | | | | | | |
| Cyclophosphamide | Immune suppressant | 50 mg/kg | IV | | | | | | | | | | | | | | | | | | | | | | | | | | | |
| Tacrolimus | Immune suppressant | 0.01 mg/kg | IM | | | | | | | | | | | | | | | | | | | | | | | | | | | |
| Fluconazole | Anti-fungal | 5 mg/kg | oral | | | | | | | | | | | | | | | | | | | | | | | | | | | |
| Blood transfusion | | | IV | | | | | | | | | | | | | | | | | | | | | | | | | | | |
| Eltrombopag | Support platelet engraftment | 1.5 mg/kg | oral | | | | | | | | | | | | | | | | | | | | | | | | | | | |
| Necropsy | | | | | | | | | | | | | | | | | | | | | | | | | | | | | | |

### MCM-D

days post-transplant

| Treatment | Purpose | Dose | Route | -20 | .. | -14 | .. | - 7 | .. | -2 | -1 | 0 | 1 | 2 | 3 | 4 | 5 | .. | 11 | .. | 18 | .. | 25 | .. | 33 | .. | 39 |
|---|---|---|---|---|---|---|---|---|---|---|---|---|---|---|---|---|---|---|---|---|---|---|---|---|---|---|---|
| ART- TDF FTC | Anti-retroviral | 20mg/kg 40mg/kg | SQ SQ | | | | | | | | | | | | | | | | | | | | | | | | |
| ART-RAL | Anti-retroviral | 100mg | oral | | | | | | | | | | | | | | | | | | | | | | | | |
| Trimethoprim/ sulfamethoxazole | Gut decontamination | 200mg 40mg | oral | | | | | | | | | | | | | | | | | | | | | | | | |
| Ceftriaxone | Antibiotic | 50 mg/kg | IM | | | | | | | | | | | | | | | | | | | | | | | | |
| Cefazolin | Antibiotic | 330 mg/ml | IM | | | | | | | | | | | | | | | | | | | | | | | | |
| Total body irradiation | Immune conditioning | 5Gy for 2 days | - | | | | | | | | | | | | | | | | | | | | | | | | |
| TCRα/β-depleted cells | | 8.6x10^8 cells | IV | | | | | | | | | | | | | | | | | | | | | | | | |
| Cyclophosphamide | Immune suppressant | 50 mg/kg | IV | | | | | | | | | | | | | | | | | | | | | | | | |
| Tacrolimus | Immune suppressant | 0.01 mg/kg | IM | | | | | | | | | | | | | | | | | | | | | | | | |
| Fluconazole | Anti-fungal | 5 mg/kg | oral | | | | | | | | | | | | | | | | | | | | | | | | |
| Blood transfusion | | | IV | | | | | | | | | | | | | | | | | | | | | | | | |
| N-acetyl L-cysteine | Support platelet engraftment | 50mg/kg | oral | | | | | | | | | | | | | | | | | | | | | | | | |
| Necropsy | | | | | | | | | | | | | | | | | | | | | | | | | | | |
