## Supplementary Table 2 for "MHC-matched allogeneic bone marrow transplants fail to eliminate SHIV-infected cells from ART-suppressed Mauritian cynomolgus macaques"

**Supplementary Table 2.** Panel of single nucleotide polymorphisms (SNPs) and primers used to identify diagnostic SNPs distinguishing donor and recipient MCMs.

| Gene | Chromosome | Position | Allele Change | Allele Specific Primers | Locus Specific Primer |
|---|---|---|---|---|---|
| CELSR2 | 1 | 112321563 | C → T | CCCTGCAGTGAATTACGA[C/T] | GCGCTTCCTGCCCTCATC |
| GIPR | 19 | 52125587 | C → T | ATGGTCATGAGGATGGG[G/A] | GCGAGCGCAACGAAGTCAA |
| GPR107 | 15 | 8694018 | C → G | CCTCAGGGTCGGCT[C/G] | GCTCCTTGTTCAGTGTGACAACAT |
| GPR139 | 20 | 19030347 | A → G | ATATCGCTGTCTGCCA[T/C] | GCGTGCTGATGTAGTCTTCAGTC |
| GPR149 | 2 | 133230027 | A → T | CGCTTTGATCCTAGCACTTAC[A/T] | GCCTTGTGAGCAACTGCACTTTTA |
| GPR183 | 17 | 79668813 | G → A | GCCTCTGCATTACAGCCT[C/T] | GCATGACAACCAAGGCTAGTAAGTTT |
| GPR68 | 7 | 154450353 | G → T | TGATGTAGATGTTCTCGTAGAG[G/T] | GCGCGTGTACCTGTGCAA |
| GPR98 | 6 | 86863653 | C → T | TATGGAAAACCAGAAGATTGAAAG[C/T] | GCCTCACATCTCCTTTAGTCCCT |
| HTR5A | 3 | 192188304 | A → G | GGTCAACTATTGGGACATAC[T/C] | GCGGATAGGAGACCATAGTTCCA |
| MC4R | 18 | 53659440 | G → A | GATTGCTGTCCTCCC[C/T] | GCCCAATCAGGATGGTCAAAGTAAT |
| P2RY11 | 19 | 9916169 | G → A | AAGCTGCGTGTGGC[G/A] | GCCCGAGCATCCACGTTGA |
| TAAR1 | 4 | 131253869 | C → T | ATCTCTTCAGCGCCTTTGAA[G/A] | GCTGTCTTTCATCTCCATTGACCG |

We genotyped each donor/recipient pair using the rhAMP SNP Assay (IDT) to identify diagnostic SNPs, with preference given to homozygous/homozygous mismatches. The chromosome positions are relative to the rhesus macaque genome assembly rheMac2.
