## Supplementary Table 3 for "MHC-matched allogeneic bone marrow transplants fail to eliminate SHIV-infected cells from ART-suppressed Mauritian cynomolgus macaques"

**Supplementary Table 3.** SNPs used in this study to distinguish donor and recipient MCMs.

| Donor | Recipient | SNP | Donor/Recipient Expected | Donor Basepair | Recipient Basepair |
|---|---|---|---|---|---|
| MCM-A | MCM-E | CELSR2 | Homozygous/Homozygous | C | T |
| MCM-B | MCM-F | CELSR2 | Homozygous/Heterozygous | C | C / T |
| | | MC4R | Heterozygous/Homozygous | C / T | T |
| MCM-C | MCM-G | MC4R | Homozygous/Homozygous | C | T |
| MCM-D | MCM-H | HTR5A | Heterozygous/Homozygous | G / A | G |
| | | GPR183 | Homozygous/Heterozygous | T | C / T |
