## Supplementary Table 4 for "MHC-matched allogeneic bone marrow transplants fail to eliminate SHIV-infected cells from ART-suppressed Mauritian cynomolgus macaques"

**Supplementary Table 4.** Sequencing primers for amplifying diagnostic SNPs.

Illumina Sequencing Primers

| Gene | 5' Primer | 3' Primer | Amplicon Length (bp) |
|---|---|---|---|
| CELSR2 | GGAGCTGCTCCTGGGTGAC | CAACACCTACCAAAGGAGCCTT | 150 |
| GPR183 | CACCTCCTCAGGGAAATGA | GAACAATGACAACCAAGGC | 132 |
| HTR5A | GAGACCATAGTTCCAGGCT | TCCCTGCTTTCATGGATAGGA | 165 |
| MC4R | TTGGCTCTCATGGCTTCTCTC | CAGCAGACAACAAAGACGCC | 161 |
